# Defining the most potent osteoinductive culture conditions for MC3T3-E1 cells reveals no implication of oxidative stress or energy metabolism

**DOI:** 10.1101/2024.02.28.582483

**Authors:** Alexandra Semicheva, Ufuk Ersoy, Aphrodite Vasilaki, Ioanna Myrtziou, Ioannis Kanakis

## Abstract

MC3T3-E1 preosteoblastic cell line is widely utilised as a reliable *in vitro* system to assess bone formation. However, the experimental growth conditions for these cells hugely diverge and, particularly, the osteogenic medium (OSM) composition varies in research studies. Therefore, we aimed to define the ideal culture conditions for MC3T3-E1 subclone 4 cells with regards to mineralization capacity and explore if oxidative stress or cellular metabolism processes are implicated. Cells were treated with 9 different combinations of long-lasting ascorbate (Asc) and β-glycerophosphate (βGP) and osteogenesis/calcification were evaluated at 3 different time-points by qPCR, Western blotting and bone nodule staining. Key molecules of oxidative and metabolic pathways were also assessed. It was found that sufficient mineral deposition was achieved only in 150μg.mL^-1^/2mM Asc/βGP combination at 21d in OSM and this was supported by *Runx2, Alpl, Bglap* and *Col1a1* expression levels increase. NOX2 and SOD2 as well as PGC1α and Tfam were also monitored as indicators of redox and metabolic processes, respectively, where no differences were observed. Elevation in OCN protein levels and ALP activity showed that mineralisation comes as a result of these differences. This work defines the most appropriate culture conditions for MC3T3-E1 cells and could be used by other research laboratories in this field.

## INTRODUCTION

Osteoblasts are specialized bone-forming cells that play a crucial role in bone development, homeostasis, and repair [1]. Understanding the factors that regulate osteoblast differentiation and mineralisation is of great importance in the field of bone tissue engineering, regenerative medicine as well as to understand the pathophysiology of common skeletal disorders like osteoporosis. *In vitro* cell culture models offer a valuable tool for studying the complex processes involved in osteogenesis and ossification, allowing researchers to investigate the effects of different factors on osteoblast behaviour and bone formation [2].

The MC3T3-E1 cell line, derived from mouse calvaria, is widely used as a model system for studying bone formation *in vitro* [3]. MC3T3-E1 cells exhibit characteristics of pre-osteoblasts and have the capacity to undergo osteogenic differentiation when cultured under appropriate conditions [4]. Several subclones of MC3T3-E1 cells have been established, each with its unique characteristics and responsiveness to differentiation stimuli [5]. One key factor that influences osteoblastic function *in vitro* is the culture medium composition. Different culture media formulations have been developed to provide optimal conditions for MC3T3-E1osteoblastic induction and calcification potential. Alpha-Minimum Essential Medium (α-MEM) supplemented with fetal bovine serum (FBS) is commonly used as a basal medium for culturing MC3T3-E1 cells [5]. FBS provides essential growth factors and nutrients to support cell proliferation and survival. However, during osteogenic differentiation, the addition of specific supplements, such as ascorbic acid (Asc) and beta-glycerophosphate (βGP), is necessary to induce the osteoblastic phenotype with mineralisation capacity [6].

Ascorbic acid, a vitamin C derivative, is a critical component in the osteogenic differentiation of MC3T3-E1 cells. It serves as a cofactor for prolyl hydroxylase, an enzyme involved in collagen synthesis, which is essential for bone matrix formation [7]. Ascorbic acid also has antioxidant properties that protect against oxidative stress, which can impair osteoblast function [8]. On the other hand, βGP, an organic phosphate source, is another crucial component in hydroxyapatite deposition on bone matrix produced by MC3T3-E1 cells. It serves as a substrate for alkaline phosphatase, an enzyme that plays a pivotal role in mineralisation by hydrolysing phosphate esters [9,10]. Supplementation of βGP enhances osteogenic commitment of MC3T3-E1 cells. Researchers have explored the effects of different concentrations of βGP on osteoblast behaviour and found that an optimal range (2-10 mM) promotes mineralisation [11-13]. However, it has been reported that concentrations above 5mM results in non-specific staining of the bone nodules being formed by mouse osteoblasts [6,14].

In the current literature, there is an inconsistency between different *in vitro* experiments about the use of specific concentrations for Asc and βGP and lack of understanding how essential mechanisms in osteoblast differentiation and mineralisation, like oxidative stress and energy metabolism, are affected by their concentration. Oxidative stress, resulting from an imbalance between reactive oxygen species (ROS) production and antioxidant defenses, plays a crucial role in osteoblast differentiation and function [15,16]. NADPH oxidase 2 (NOX2), a key enzyme in reactive oxygen species (ROS) production, has a multifaceted role in osteoblasts. It is primarily involved in redox signaling, regulating osteoblast differentiation, and bone homeostasis [17]. NOX2-derived ROS act as secondary messengers, participating in intracellular signaling cascades crucial for osteoblast proliferation and differentiation, but excessive ROS production can lead to cellular damage and impaired bone formation [18]. Moreover, NOX2 activity is linked to the modulation of osteoclastogenesis, thus influencing bone remodeling [19]. Another important enzyme, superoxide dismutase 2 (SOD2) operates as a critical defender against oxidative stress, counteracting the harmful effects of increased ROS [20]. SOD2 deficiency can lead to mitochondrial dysfunction and increased ROS levels, disrupting osteoblast function and potentially leading to a compromised bone microenvironment [21].

Energy metabolism is tightly linked to osteoblast activity, as these cells demand substantial energy for the synthesis of bone matrix components. Proper mitochondrial function is critical for osteoblast differentiation and mineralisation while dysregulation of mitochondrial activity can impair osteoblast function, affecting bone formation [22]. Peroxisome Proliferator-Activated Receptor-Gamma Coactivator 1 alpha (PGC1α) is a transcriptional coactivator that regulates energy metabolism and mitochondrial biogenesis [23]. It has been reported that PGC-1α enhances the generation of new mitochondria, supporting the energetic demands of osteoblasts and facilitating their differentiation into mature bone-forming cells [24] and depletion of PGC1α in osteoblasts contributes to decreased bone mass in osteoporosis [25]. Furthermore, mitochondrial transcription factor A (Tfam) is a nuclear-encoded protein primarily known for its role in regulating mitochondrial DNA maintenance [26]. It has been reported that deletion of Tfam leads to mitochondrial dysfunction of osteoblasts in Tfam-knockout mice and subsequent decreased bone formation [27].

Our aim was to define the ideal culture conditions for MC3T3-E1 preosteoblasts in terms of mineralising capacity induced by stable chemical forms for Asc and βGP. For this reason, we selected subclone 4 which has been shown to exhibit a higher calcification rate than other subclones [28]. We also investigated potential pathways that can influence this process and explored if oxidative stress-related molecules and mitochondrial activity can be influenced by different culturing conditions.

## MATERIALS AND METHODS

### Cell culture

MC3T3-E1 subclone 4 (ATCC CRL-2593) preosteoblastic cell line was obtained from the American Tissue Culture Collection (Rockville, MD). Cells were cultured in alpha-MEM with Glutamax™ (Gibco, UK) and nucleosides, containing 10% heat-inactivated FBS and penicillin (100 IU/ml)/streptomycin (100 μg/ml) (Invitrogen) in a humidified 5% CO2 incubator at 37°C, as previously described [29,30]. Upon reaching confluence, cells were harvested using trypsin/EDTA (Gibco, UK) and seeded onto 6-well plates (10^5^cells/well). All wells contained cells from the second passage. Two days after seeding and adherence of the cells, 2ml of OSM containing different concentrations of Asc and βGP (Sigma, UK) were added to the wells. The actual form for these OSM factors were: Asc - L-Ascorbic acid 2-phosphate sesquimagnesium salt hydrate (Sigma, A8960) and βGP - β-Glycerophosphate disodium salt hydrate (Sigma, G9422). The combinations of Asc/βGP concentrations were (μg/mL Asc/mM βGP): 100/2, 100/3, 100/4, 150/2. 150/3, 150/4, 200/2, 200/3 and 200/4 (9 experimental conditions). Cells without OSM (undifferentiated, UN) served as control. All conditions were assessed in triplicate and in all wells conditioned medium was changed every 3 days with freshly prepared OSM. Three sets of time-points were used for analysis: 7d and 14d for gene expression by qPCR, ALP activity by spectrophotometry and protein abundance by Western blot and 21d for terminal mineralisation evaluation.

### Bone nodule staining

Mineralisation capacity was assessed by ARS (Sigma, UK) staining. In brief, on the 21st day of culture in OSM, wells were washed twice with PBS (without Ca_2+_) and cells were fixed with 1mL of 10% neutral buffered formalin (NBF) for 20min. Following three additional washes with PBS, 1mL of 4mM ARS (pH 4.2) was added for 30min and wells were extensively washed with distilled H_2_O until no colour could be observed in the washes. Wells were photographed and 10x pictures were taken with Leica DM IL LED (Leica, UK) inverted brightfield microscope. Bone nodule surface area was calculated using ImageJ (NIH), as previously described [29,30].

### Alkaline phosphatase (ALP) activity assay

ALP activity assay in cell culture supernatants was conducted following a 7d and 14d period of culture with OSM. Twenty-four hours before supernatant collection, culture medium was replaced αMEM without phenol (Gibco, UK) OSM to avoid interference with the colorimetric method. Using 96-well plates, equal volumes (50μL) of p-nitrophenyl phosphate (p-NPP) in 0.01M glycine buffer (pH 10.5) solution was added to 100μL of supernatants (in duplicate), followed by incubation for 60min at 37°C. The reaction was terminated by adding 20μL of 1N NaOH. The enzyme activity of p-NPP hydrolysis was measured with a microplate reader at 405 nm, and p-nitrophenol (p-NP) was used to construct the standard curve. ALP activity was expressed as Units/L.

### Quantitative polymerase chain reaction (qPCR)

To compare gene expression levels, total RNA was isolated from MC3T3-E1 cells of all conditions from 3 biological replicates at 7d and 14d of culture using TRiZOL reagent (Invitrogen, CA, USA), cleaned-up using the RNeasy kit (Qiagen, UK) and cDNA was synthesized with the High Capacity cDNA Transcription kit (Applied Biosystems, UK). Expression levels were measured for alkaline phosphatase (*Alpl*), collagen type 1 (*Col1a1*), runt-related transcription factor 2 (*Runx2*), bone gamma-carboxyglutamic acid-containing protein (*Bglap*), peroxisome proliferator-activated receptor gamma coactivator 1-alpha (*Pgc1α*), transcription Factor A mitochondrial (*Tfam*) and superoxide dismutase (*Sod2*) (Table S1). Real time qPCR was performed using a RotorGene 6000 (Corbett Research) instrument with SYBR (Bioline, UK) and results were analysed using beta-actin (*Actb*) as a stable reference gene for murine osteoblasts. Results were analysed using the delta delta CT method (2^-ΔΔCt^) and presented as fold change as compared to the reference gene expression levels [31].

### Western blotting

Along with RNA isolation, total protein was extracted from MC3T3-E1 cells after 7d and 14d in OSM under all experimental conditions. Cells were lysed with RIPA Lysis Buffer (Sigma, Poole, UK) containing Pierce Protease Inhibitors (ThermoFisher, UK), and the samples were homogenised by sonication for 30 seconds twice, and then centrifuged for 10 min at 14,000 x g at 4°C. The supernatant was collected and total protein content determined with a BCA Protein Assay (Sigma, Poole, UK). A total of 20μg protein extract for each sample was separated by electrophoresis on Nu-PAGE™ 4-12%, Bis-Tris Mini Protein Gel (Invitrogen, Renfrewshire, UK) and transferred to a polyvinylidene fluoride (PVDF) membrane. Actin β (ACTB) was used as loading control. Membranes were blocked with 5% bovine serum albumin (BSA) (Sigma, Poole, UK) in tris-buffered saline (TBS) for 1h and probed with primary antibodies overnight at 4°C. After incubating with a fluorescence secondary antibody (Licor rabbit anti-mouse, 1:10,000), blots were imaged on Licor Odyssey CLx (Licor, Bad Homburg, Germany) and band densities were analysed using ImageJ. Band quantifications were normalised to the reference protein.

The primary antibodies are as follows: anti-NOX2/gp91phox (ab129068, dilution 1:400), anti-ACTB (ab8227, 1:2000), anti-OCN (ab93876, 1:500) from Abcam (Cambridge, UK); anti-SOD2 (ADI-SOD-111, 1:3000) from Enzo Life Sciences (Farmingdale, New York, USA); anti-PGC1α (NBP1-04676, 1:400) from Novus Biologicals (Abingdon, UK).

### Statistical analysis

All data were analyzed with GraphPad Prism 9 software and expressed as the mean ± SD. Data sets were tested for Gaussian distribution with the D’Agostino-Pearson normality test. Comparisons between experimental conditions were performed by one-way analysis of variance (ANOVA) followed by Tukey’s multiple comparisons post hoc test. In all cases, *p* values less than 0.05 were considered statistically significant.

## RESULTS

### Bone nodules are abundantly formed under specific culture conditions

To test our hypothesis, MC3T3-E1 subclone 4 preosteoblasts were cultured in osteogenic medium (OSM) using 9 different combinations of Asc and βGP concentrations for 21d to examine the mineralisation capacity. Chemical derivatives for Asc and βGP that are stable and long acting in cell culture conditions for both factors were used (see Methods). Although, a concentration of 50 μg.mL^-1^/5mM Asc/βGP is very effective based on our previous publications with primary mouse osteoblasts [29,30], a pilot experiment showed no sign of mineralisation with the MC3T3-E1 subclone 4 cell line with the aforementioned concentrations (data not shown). Therefore, we selected the 100, 150 and 200 μg/mL of Asc for the subsequent tests and also screened a combination of 2, 3 and 4 mM βGP as 2mM is suggested to be very potent in bony structure formation in vitro [14].

Alizarin Red S (ARS) staining of mineralised bone matrix at 21d showed a remarkable statistically significant difference between the 150/2 condition as compared to the undifferentiated control, as expected, as well as other conditions (Fig. 1A-J). The 100/2 treatment also demonstrated some limited mineral deposition as compared to 150/2 but substantially higher than other conditions (Fig. 1K).

**Figure 1.**
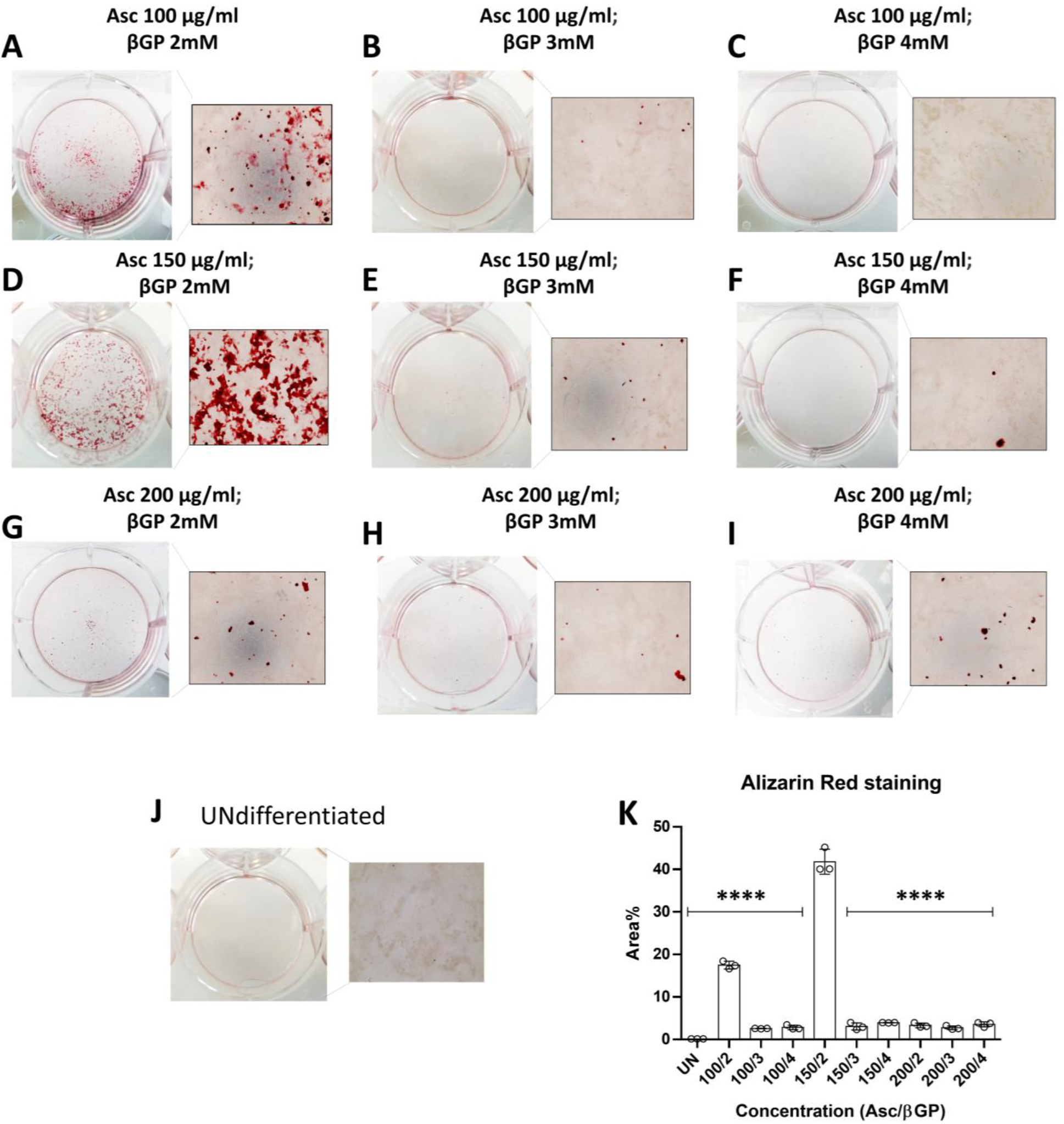
Measurement of calcified area. The effect of different combinations of Asc/βGP in OSM were measured in 6-well plates stained by ARS (A-I) in triplicate (n=3). Undifferentiated MC3T3-E1 cells (UN) were used as control (J). Quantification of stained areas (K) is presented as mean±SD. \*\*\*\**p*<0.0001 as compared to 150/2 condition.

### Asc/βGP 150/2 induces osteogenic marker gene expression and increased OCN levels

We then explored the early osteoblastic differentiation (7d) and the progression of bone matrix formation (14d) events of MC3T3-E1 subclone 4 cells in OSM. Therefore, the gene expression levels of the transcription factor *Runx2*, a master regulator of osteoblastogenesis, were measured along with *Alpl* and *Bglap*, as mineralisation markers, and *Col1a1* as bone matrix production marker at 7d and 14d of culture in OSM.

It was found that *Alpl, Col1a1* and *Bglap* genes expression levels were significantly higher in the 150/2 group as compared to the other conditions at both time-points (Fig. 2B-D). It could be also noted that for the 150/2 condition there was an increase in gene expression between 7 and 14d showing accelerating bone formation. However, *Runx2* expression did not differ between groups (Fig. 2A), except for conditions 100/4, 150/4 and 200/4 but only at 14d, suggesting that preosteoblastic cells proceed similarly to osteoblastic differentiation in a variety of Asc/βGP concentrations but respond differently regarding bone matrix secretion and mineralisation. In addition, Western blot analysis showed that OCN protein levels, which plays crucial roles in the mineralisation process, were significantly increased in the 100/2, 100/3 and 100/4 groups as compared to the 150/2 at 7d (Fig. 2E and F) but this was reversed at the 14d time-point where OCN under 150/2 treatment was found statistically elevated as compared to the other conditions, except 150/3 and 150/4 (Fig. 2G and H) suggesting that possibly this Asc concentration induces OCN production.

**Figure 2.**
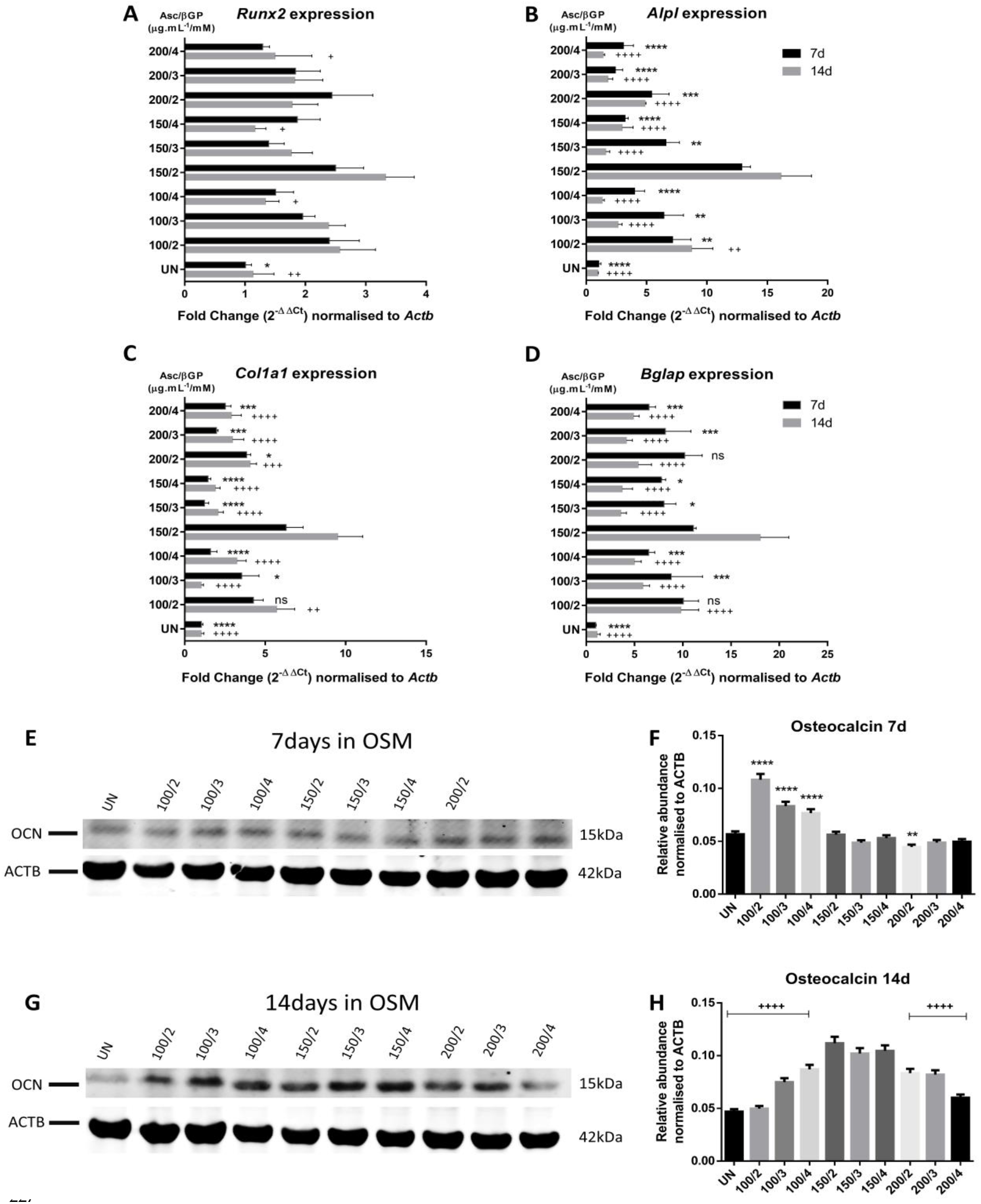
Gene/protein expression of key bone genes at 7d and 14d in OSM. *Runx2, Alpl, Bglap* and *Col1a1* gene expression (A-D) was measured by qPCR and expressed as fold change normalised to *Actb*. All comparisons were performed at 7d and 14d by ANOVA (n=3). Bands intensity in Western blots for OCN at 7d (E) and 14d (G) were quantified using ACTB protein for nomalisation and expressed as relative abundance (n=3). OCN protein abundance of all experimental conditions was compared to 150/2 group at 7d (F) and 14d (H). All data are presented as mean ± SD. ns: not significant; ^*^*p*<0.05, ^**^*p*<0.01, ^***^*p*<0.001, ^****^*p*<0.0001. Asterisks indicate comparisons versus 150/2 at 7d and crosses at 14d.

### Culture medium composition controls bone matrix calcification progress

Furthermore, microscopic imaging revealed that under all conditions bone matrix was abundantly secreted at the end stage of 21d (Fig. 3A-I) but, as expected, this was less profound in the undifferentiated cells (Fig. 3J). However, ARS staining revealed that extended mineral deposition in bone matrix nodules could be observed only in the 100/2 and 150/2 conditions (Fig. 2 A and D) which was clearly more intense in the 150/2 group.

**Figure 3.**
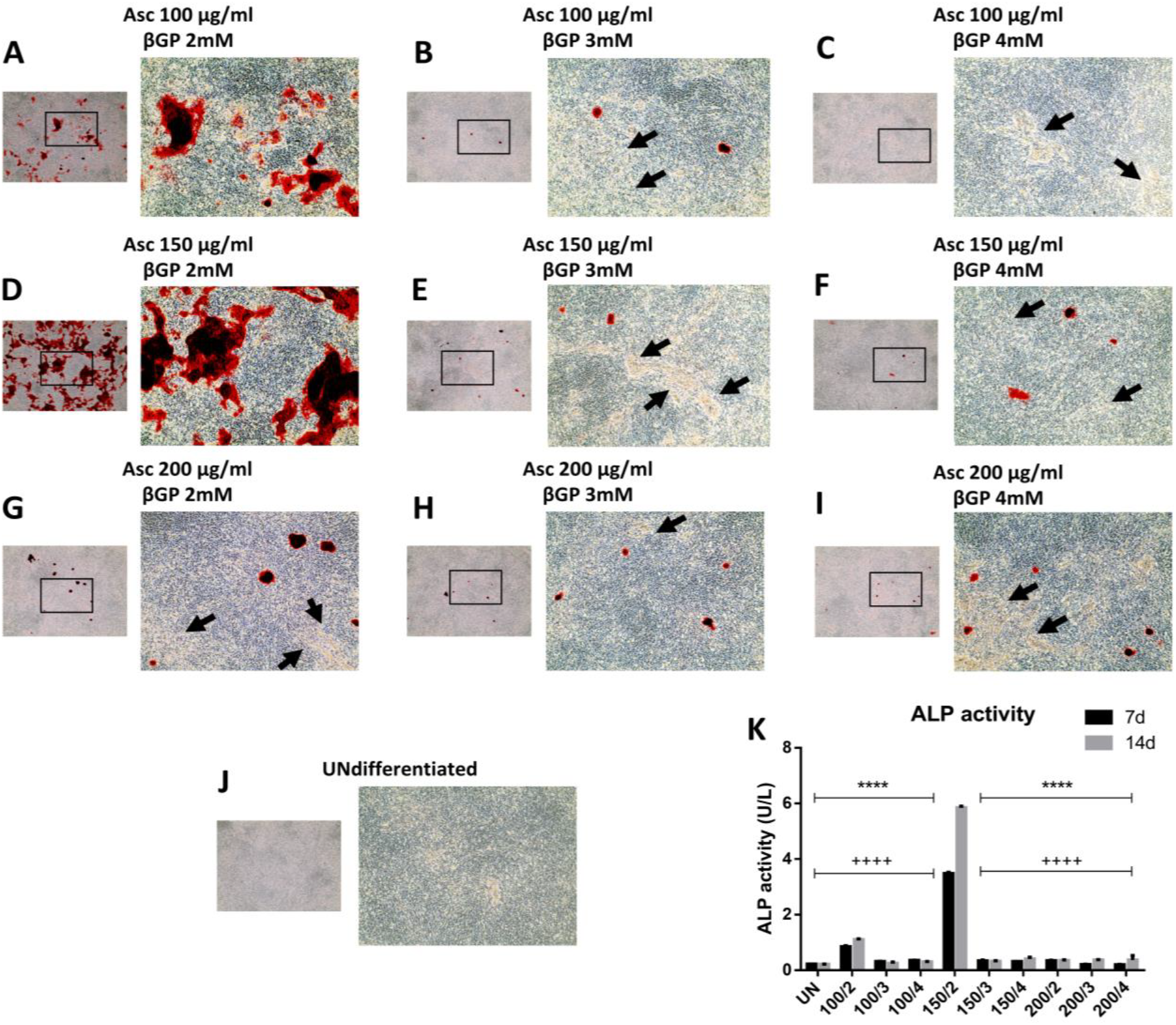
Observation of unmineralized matrix. Microscopic evaluation of bone matrix production was performed in all experimental conditions (A-J). Arrows indicate representative regions were bone matrix remains unmineralised. ALP activity (n=3) was quantified at 7d and 14d (K) and was expressed as U/L. Data are presented as mean ± SD. ^****^*p*<0.0001. Asterisks indicate comparisons versus 150/2 at 7d and crosses at 14d.

This finding suggests that in all the other conditions, except 100/2 and 150/2, matrix remains unmineralised or this process is delayed. Finally, alkaline phosphatase (ALP) activity in the supernatant of the cell cultures at 21d clearly indicates that the mineralisation process is more intense in the 150/2 wells followed by the 100/2 condition as reflected by the corresponding levels of ALP enzymatic activity (Fig. 3K), while in the other conditions it seems that there is a delay in the hydroxyapatite crystal formation and deposition.

### Compromised bone formation does not implicate oxidative stress and mitochondrial respiration

Based on our findings, we next aimed to explore whether basic processes, like energy metabolism and oxidative stress, that regulate cellular functions of osteoblasts are implicated in the altered osteoblasts’ responses under different experimental conditions.

We, therefore, measured the gene expression levels of *Sod2* and protein levels of SOD2 and NOX2 which reflect oxidative stress events, and gene expression levels of *Pgc1α* and *Tfam* as well as the protein abundance of PGC1A, as indicators of mitochondrial biogenesis and cellular energy metabolism. No statistically significant dereferences were observed for *Pgc1α* and *Tfam* at 7d or 14d for the different culture conditions (Fig. 4A-C) whereas *Sod2* expression levels for all conditions were statistically different to UN. However, it could be noted that for the majority of concentrations, expression levels of the genes were decreased at 14d as compared to the 7d and this was more profound for *Pgc1α* (Fig. 4A). Similar results were obtained from Western blots for PGC1A, SOD2 and NOX2 protein levels for the 7d (Fig. 4D-G) and 14d (Fig. H-K) in OSM. Statistically significant differences were found in different conditions and timepoints, such as for PGC1A for UN, 150/3 and 150/4 conditions, NOX2 for UN and SOD2 for 200/4 at 7d as well as for PGC1A for 100/2, 200/2, 200/3 and 200/4 conditions at 14d, but not a specific pattern could be observed.

**Figure 4.**
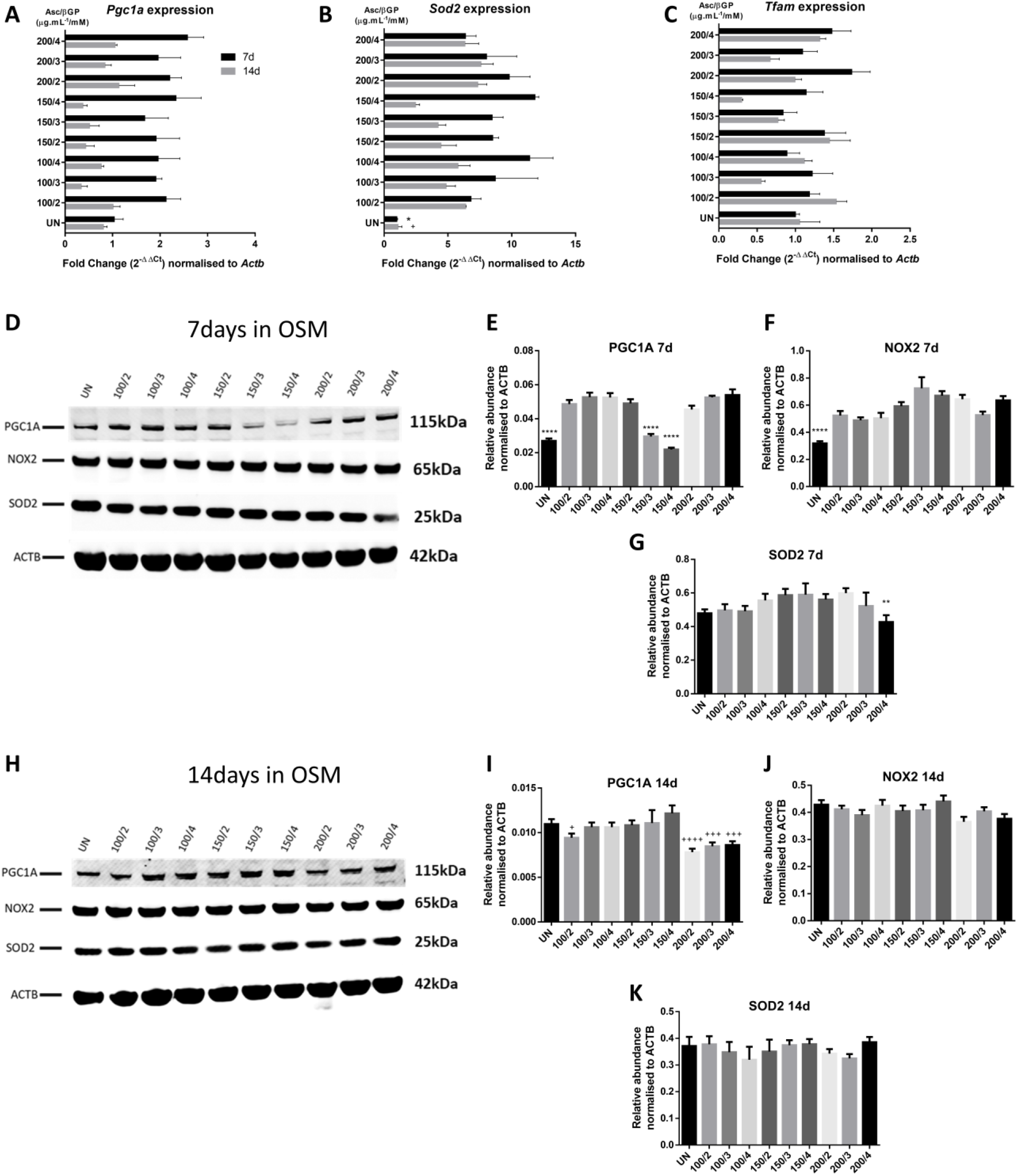
Gene/protein expression of key molecules implicated in oxidative stress and energy metabolism at 7d and 14d in OSM. *Pgc1α, Sod2* and *Tfam* gene expression (A-C) was measured by qPCR and expressed as fold change normalized to *Actb*. All comparisons were performed at 7d and 14d by ANOVA (n=3). Bands intensity in Western blots for PGC1A, NOX2and SOD2 at 7d (E-G) and 14d (H-K) were quantified using ACTB protein for nomalisation and expressed as relative abundance as compared to 150/2 group (n=3). All data are presented as mean ± SD. ns: not significant; ^*^*p*<0.05, ^**^*p*<0.01, ^***^*p*<0.001, ^****^*p*<0.0001. Asterisks indicate comparisons versus 150/2 at 7d and crosses at 14d.

## DISCUSSION

*In vitro* cell model systems resemble the physiological events of osteogenesis and calcification, and their use has been useful in the preclinical evaluation of therapeutic agents targeting bone diseases. Among cells with osteogenic potential, MC3T3-E1 cells is the most established model and different subclones have been isolated and characterised [32]. However, and although these cells are commercially available with numerous publications using this cell line, there is often lack of information regarding the use of a specific subclone in published work or consistency in the osteoinductive culture media used. Therefore, in this work, we intended to identify the most potent osteogenic culturing conditions for MC3T3-E1 subclone 4 cells, which has been shown to exhibit good osteogenic potential as compared to other sub-clones [12,32]. We report that OSM containing Asc/βGP 150μg.mL^-1^/2mM induces cells to produce abundant mineralised bone nodules at 21d, where alternative combinations failed to achieve this and were limited to secretion of uncalcified bone matrix. In addition, the production of calcified matrix under this condition is accompanied by significant differences in key osteogenic gene markers in comparison with other Asc/βGP combinations. Finally, there was no clear implication of oxidative stress and energy metabolism in this procedure as revealed by analysis of key factors involved in these processes.

Asc and βGP are supplements that are being used to induce commitment for differentiation into the osteoblastic lineage as well as to promote bone-forming capacity [12,32] since spontaneous mineralisation is not observed in *in vitro* systems [33,34]. Although the function of Asc is well characterised, mainly participating in procollagen synthesis and mature collagen assembly as a crucial cofactor of propyl/lysyl hydroxylases [35-37], the exact role of βGP has not been fully uncovered besides being a substrate for hydrolysis by alkaline phosphatase. Furthermore, βGP concentrations used in the literature vary between 2-10mM depending on the cell type (cell lines, primary, bone marrow mesenchymal stem cells etc.) [28,38]. On the other hand, it has been reported that βGP addition above 5mM can be cytotoxic and result in nucleation of non-bone-specific minerals that could lead to overestimation of mineralisation capacity [9,14]. We used in our experimental setting a concentration range of 2-4mM βGP and found that the 2mM is more efficient in bone nodule formation by MC3T3-E1 subclone 4 cells, consistent with Chung et al. date? who showed that higher concentrations can lead to formation of calcium-phosphate crystals other than hydroxyapatite [39]. Therefore, it is crucial to use a low βGP concentration when assessing mineralisation in order to ensure a real effect and avoid unspecific widespread calcification. Furthermore, Asc seems to be utilised in a wide range of concentrations as well, but in most cases conventional Asc is added to the OSM at a range of 50-100μg/mL [40-43]. We urged to explore the effect of the long-lasting L-Ascorbic acid 2-phosphate in our culture system and found that 150μg/mL resulted in the formation of specific and abundant bone nodules.

In addition, we report that this Asc form can induce osteogenic markers gene expression in agreement with previous reports [44,45]. However, this was more evident and statistically significant in the 150/2 condition as compared to the other experimental conditions and consistent throughout the time course of the experimental period. *Col1a1* gene expression analysis showed a significant and persistent increase in 150/2 group at both 7d and 14d time-points, forming a mature bone ECM which is subsequently being calcified [46]. Notably, cells in alternative conditions secreted bone matrix as well at 21d in OSM, but these nodules remained unmineralised. Similarly, *Alpl* and *Bglap* gene levels were elevated in 150/2 while a decrease was observed in all other conditions, except the 100/2 which also showed a degree of mineralisation, highlighting that βGP at 2mM is the most potent osteoinductive condition. In contrast, OCN protein showed an increase at 150/2 combination only at 14d. Finally, *Runx2* levels did not differ among the experimental conditions showing that MC3T3-E1 subclone 4 preosteoblastic cells can be differentiated and form mature osteoblasts at various combinations of Asc/βGP. Collectively, our results indicate that the differences observed between the nine Asc/βGP combinations used are caused by the alterations of the mineralisation capacity, as a result of alkaline phosphatase and osteocalcin activity changes, rather than a phenotypic modification, showing that their role is crucial [47,48]. It is also important to note that the reported changes and the delay in biomineralization refer to the specific experimental time course of 21d in OSM.

In an effort to further explore the cause in the aforementioned results, we evaluated key molecules of oxidative stress and energy metabolism, cellular processes that affect bone health and bone-related diseases such as osteoporosis [16,49-54]. We first examined the gene and/or protein expression levels of Pgc1a and Tfam, as important factors of mitochondrial biogenesis in bone cells [24]. It is known that the AMPK/PGC1α axis is activated during osteoblast differentiation in C3H10T1/2 and Tfam is also up-regulated in osteoblast-like MG63 osteosarcoma cells [55,56]. On the other hand, osteoblast-specific Sod2 deficient mice develop an osteoporotic phenotype [21], while Nox2 increase leads to osteocyte apoptosis [57] and glucocorticoid-induced pre-osteoblast death [58](41). However, we did not find any differences between our experimental conditions, a result which supports our main conclusion that the mineralisation process is the key difference in the current study.

## Conclusions

In conclusion, here we define the most potent osteogenic culture conditions in which MC3T3-E1 subclone 4 preosteoblastic cells differentiate into mature osteoblasts and produce bone matrix which is mineralised resulting in a sufficient number of bone nodules quantifiable by ARS staining at 21d. This OSM is specifically supplemented with 150μg/mL of L-Ascorbic acid 2-phosphate sesquimagnesium salt hydrate and 2mM of β-Glycerophosphate disodium salt hydrate. In addition, this finding is supported by a significant increase in osteogenic markers and ALP and OCN proteins which are key regulators of calcification. In combination with the gene/protein expression results of Pgc1a, Tfam, Sod2 and Nox2, we conclude that the final outcome in bone formation is due to the effect of the specific OSM on mineralization with no implication of oxidative stress or mitochondrial biogenesis processes. Our results could be used as a standard protocol from other research teams and could assist in obtaining reliable and repeatable results in preclinical testing of osteoanabolic agents without under- or overestimation of the mineralisation outcome.

## Author Contributions

I.K. and I.M.K. designed the experiments with input from U.E. and A.V.; A.S., U.E., I.K. and I.M.K. performed the experiments and acquired and analysed the data; I.K. and I.M.K. drafted the manuscript, which was critically revised and approved by all co-authors. All authors have read and agreed to the published version of the manuscript.

## Institutional Review Board Statement

Not applicable

## Informed Consent Statement

Not applicable

## Data Availability Statement

Not applicable

## Acknowledgments

We acknowledge the support of the technical team of the Chester Medical School, University of Chester, UK.

## References

1. Dirckx, N.; Moorer, M.C.; Clemens, T.L.; Riddle, R.C. The role of osteoblasts in energy homeostasis. Nat Rev Endocrinol 2019, 15, 651–665, doi:10.1038/s41574-019-0246-y.

2. Czekanska, E.M.; Stoddart, M.J.; Richards, R.G.; Hayes, J.S. In search of an osteoblast cell model for in vitro research. Eur Cell Mater 2012, 24, 1–17, doi:10.22203/ecm.v024a01.

3. Sudo, H.; Kodama, H.A.; Amagai, Y.; Yamamoto, S.; Kasai, S. In vitro differentiation and calcification in a new clonal osteogenic cell line derived from newborn mouse calvaria. J Cell Biol 1983, 96, 191–198, doi:10.1083/jcb.96.1.191.

4. Kurihara, N.; Ishizuka, S.; Kiyoki, M.; Haketa, Y.; Ikeda, K.; Kumegawa, M. Effects of 1,25-dihydroxyvitamin D3 on osteoblastic MC3T3-E1 cells. Endocrinology 1986, 118, 940–947, doi:10.1210/endo-118-3-940.

5. Wang, D.; Christensen, K.; Chawla, K.; Xiao, G.; Krebsbach, P.H.; Franceschi, R.T. Isolation and characterization of MC3T3-E1 preosteoblast subclones with distinct in vitro and in vivo differentiation/mineralization potential. J Bone Miner Res 1999, 14, 893–903, doi:10.1359/jbmr.1999.14.6.893.

6. Taylor, S.E.; Shah, M.; Orriss, I.R. Generation of rodent and human osteoblasts. Bonekey Rep 2014, 3, 585, doi:10.1038/bonekey.2014.80.

7. Rappu, P.; Salo, A.M.; Myllyharju, J.; Heino, J. Role of prolyl hydroxylation in the molecular interactions of collagens. Essays Biochem 2019, 63, 325–335, doi:10.1042/EBC20180053.

8. Aghajanian, P.; Hall, S.; Wongworawat, M.D.; Mohan, S. The Roles and Mechanisms of Actions of Vitamin C in Bone: New Developments. J Bone Miner Res 2015, 30, 1945–1955, doi:10.1002/jbmr.2709.

9. Orriss, I.R.; Utting, J.C.; Brandao-Burch, A.; Colston, K.; Grubb, B.R.; Burnstock, G.; Arnett, T.R. Extracellular nucleotides block bone mineralization in vitro: evidence for dual inhibitory mechanisms involving both P2Y2 receptors and pyrophosphate. Endocrinology 2007, 148, 4208–4216, doi:10.1210/en.2007-0066.

10. Bellows, C.G.; Aubin, J.E.; Heersche, J.N. Initiation and progression of mineralization of bone nodules formed in vitro: the role of alkaline phosphatase and organic phosphate. Bone Miner 1991, 14, 27–40, doi:10.1016/0169-6009(91)90100-e.

11. Fratzl-Zelman, N.; Fratzl, P.; Horandner, H.; Grabner, B.; Varga, F.; Ellinger, A.; Klaushofer, K. Matrix mineralization in MC3T3-E1 cell cultures initiated by beta-glycerophosphate pulse. Bone 1998, 23, 511–520, doi:10.1016/s8756-3282(98)00139-2.

12. Quarles, L.D.; Yohay, D.A.; Lever, L.W.; Caton, R.; Wenstrup, R.J. Distinct proliferative and differentiated stages of murine MC3T3-E1 cells in culture: an in vitro model of osteoblast development. J Bone Miner Res 1992, 7, 683–692, doi:10.1002/jbmr.5650070613.

13. Wenstrup, R.J.; Fowlkes, J.L.; Witte, D.P.; Florer, J.B. Discordant expression of osteoblast markers in MC3T3-E1 cells that synthesize a high turnover matrix. J Biol Chem 1996, 271, 10271–10276, doi:10.1074/jbc.271.17.10271.

14. Orriss, I.R.; Taylor, S.E.; Arnett, T.R. Rat osteoblast cultures. Methods Mol Biol 2012, 816, 31–41, doi:10.1007/978-1-61779-415-5_3.

15. Zhu, C.; Shen, S.; Zhang, S.; Huang, M.; Zhang, L.; Chen, X. Autophagy in Bone Remodeling: A Regulator of Oxidative Stress. Front Endocrinol (Lausanne) 2022, 13, 898634, doi:10.3389/fendo.2022.898634.

16. Marcucci, G.; Domazetovic, V.; Nediani, C.; Ruzzolini, J.; Favre, C.; Brandi, M.L. Oxidative Stress and Natural Antioxidants in Osteoporosis: Novel Preventive and Therapeutic Approaches. Antioxidants (Basel) 2023, 12, doi:10.3390/antiox12020373.

17. Schroder, K. NADPH oxidases in bone homeostasis and osteoporosis. Free Radic Biol Med 2019, 132, 67–72, doi:10.1016/j.freeradbiomed.2018.08.036.

18. Tian, Y.; Ma, X.; Yang, C.; Su, P.; Yin, C.; Qian, A.R. The Impact of Oxidative Stress on the Bone System in Response to the Space Special Environment. Int J Mol Sci 2017, 18, doi:10.3390/ijms18102132.

19. Sasaki, H.; Yamamoto, H.; Tominaga, K.; Masuda, K.; Kawai, T.; Teshima-Kondo, S.; Rokutan, K. NADPH oxidase-derived reactive oxygen species are essential for differentiation of a mouse macrophage cell line (RAW264.7) into osteoclasts. J Med Invest 2009, 56, 33–41, doi:10.2152/jmi.56.33.

20. Gao, J.; Feng, Z.; Wang, X.; Zeng, M.; Liu, J.; Han, S.; Xu, J.; Chen, L.; Cao, K.; Long, J., et al. SIRT3/SOD2 maintains osteoblast differentiation and bone formation by regulating mitochondrial stress. Cell Death Differ 2018, 25, 229–240, doi:10.1038/cdd.2017.144.

21. Schoppa, A.M.; Chen, X.; Ramge, J.M.; Vikman, A.; Fischer, V.; Haffner-Luntzer, M.; Riegger, J.; Tuckermann, J.; Scharffetter-Kochanek, K.; Ignatius, A. Osteoblast lineage Sod2 deficiency leads to an osteoporosis-like phenotype in mice. Dis Model Mech 2022, 15, doi:10.1242/dmm.049392.

22. Dobson, P.F.; Dennis, E.P.; Hipps, D.; Reeve, A.; Laude, A.; Bradshaw, C.; Stamp, C.; Smith, A.; Deehan, D.J.; Turnbull, D.M., et al. Mitochondrial dysfunction impairs osteogenesis, increases osteoclast activity, and accelerates age related bone loss. Sci Rep 2020, 10, 11643, doi:10.1038/s41598-020-68566-2.

23. Zhang, Y.; Castellani, L.W.; Sinal, C.J.; Gonzalez, F.J.; Edwards, P.A. Peroxisome proliferator-activated receptor-gamma coactivator 1alpha (PGC-1alpha) regulates triglyceride metabolism by activation of the nuclear receptor FXR. Genes Dev 2004, 18, 157–169, doi:10.1101/gad.1138104.

24. Buccoliero, C.; Dicarlo, M.; Pignataro, P.; Gaccione, F.; Colucci, S.; Colaianni, G.; Grano, M. The Novel Role of PGC1alpha in Bone Metabolism. Int J Mol Sci 2021, 22, doi:10.3390/ijms22094670.

25. Yu, B.; Huo, L.; Liu, Y.; Deng, P.; Szymanski, J.; Li, J.; Luo, X.; Hong, C.; Lin, J.; Wang, C.Y. PGC-1alpha Controls Skeletal Stem Cell Fate and Bone-Fat Balance in Osteoporosis and Skeletal Aging by Inducing TAZ. Cell Stem Cell 2018, 23, 615–623, doi:10.1016/j.stem.2018.09.001.

26. Larsson, N.G.; Wang, J.; Wilhelmsson, H.; Oldfors, A.; Rustin, P.; Lewandoski, M.; Barsh, G.S.; Clayton, D.A. Mitochondrial transcription factor A is necessary for mtDNA maintenance and embryogenesis in mice. Nat Genet 1998, 18, 231–236, doi:10.1038/ng0398-231.

27. Yoshioka, H.; Komura, S.; Kuramitsu, N.; Goto, A.; Hasegawa, T.; Amizuka, N.; Ishimoto, T.; Ozasa, R.; Nakano, T.; Imai, Y., et al. Deletion of Tfam in Prx1-Cre expressing limb mesenchyme results in spontaneous bone fractures. J Bone Miner Metab 2022, 40, 839–852, doi:10.1007/s00774-022-01354-2.

28. Hwang, P.W.; Horton, J.A. Variable osteogenic performance of MC3T3-E1 subclones impacts their utility as models of osteoblast biology. Sci Rep 2019, 9, 8299, doi:10.1038/s41598-019-44575-8.

29. Kanakis, I.; Liu, K.; Poulet, B.; Javaheri, B.; van ‘t Hof, R.J.; Pitsillides, A.A.; Bou-Gharios, G. Targeted Inhibition of Aggrecanases Prevents Articular Cartilage Degradation and Augments Bone Mass in the STR/Ort Mouse Model of Spontaneous Osteoarthritis. Arthritis Rheumatol 2019, 71, 571–582, doi:10.1002/art.40765.

30. Kanakis, I.; Alameddine, M.; Scalabrin, M.; van ‘t Hof, R.J.; Liloglou, T.; Ozanne, S.E.; Goljanek-Whysall, K.; Vasilaki, A. Low protein intake during reproduction compromises the recovery of lactation-induced bone loss in female mouse dams without affecting skeletal muscles. FASEB J 2020, 34, 11844–11859, doi:10.1096/fj.202001131R.

31. Livak, K.J.; Schmittgen, T.D. Analysis of relative gene expression data using real-time quantitative PCR and the 2(-Delta Delta C(T)) Method. Methods 2001, 25, 402–408, doi:10.1006/meth.2001.1262.

32. Barros, N.M.; Hoac, B.; Neves, R.L.; Addison, W.N.; Assis, D.M.; Murshed, M.; Carmona, A.K.; McKee, M.D. Proteolytic processing of osteopontin by PHEX and accumulation of osteopontin fragments in Hyp mouse bone, the murine model of X-linked hypophosphatemia. J Bone Miner Res 2013, 28, 688–699, doi:10.1002/jbmr.1766.

33. Chang, D.J.; Ji, C.; Kim, K.K.; Casinghino, S.; McCarthy, T.L.; Centrella, M. Reduction in transforming growth factor beta receptor I expression and transcription factor CBFa1 on bone cells by glucocorticoid. J Biol Chem 1998, 273, 4892–4896, doi:10.1074/jbc.273.9.4892.

34. Xiao, G.; Cui, Y.; Ducy, P.; Karsenty, G.; Franceschi, R.T. Ascorbic acid-dependent activation of the osteocalcin promoter in MC3T3-E1 preosteoblasts: requirement for collagen matrix synthesis and the presence of an intact OSE2 sequence. Mol Endocrinol 1997, 11, 1103–1113, doi:10.1210/mend.11.8.9955.

35. Murad, S.; Grove, D.; Lindberg, K.A.; Reynolds, G.; Sivarajah, A.; Pinnell, S.R. Regulation of collagen synthesis by ascorbic acid. Proc Natl Acad Sci U S A 1981, 78, 2879–2882, doi:10.1073/pnas.78.5.2879.

36. Peterkofsky, B. Ascorbate requirement for hydroxylation and secretion of procollagen: relationship to inhibition of collagen synthesis in scurvy. Am J Clin Nutr 1991, 54, 1135S–1140S, doi:10.1093/ajcn/54.6.1135s.

37. Pihlajaniemi, T.; Myllyla, R.; Kivirikko, K.I. Prolyl 4-hydroxylase and its role in collagen synthesis. J Hepatol 1991, 13 Suppl 3, S2–7, doi:10.1016/0168-8278(91)90002-s.

38. Izumiya, M.; Haniu, M.; Ueda, K.; Ishida, H.; Ma, C.; Ideta, H.; Sobajima, A.; Ueshiba, K.; Uemura, T.; Saito, N., et al. Evaluation of MC3T3-E1 Cell Osteogenesis in Different Cell Culture Media. Int J Mol Sci 2021, 22, doi:10.3390/ijms22147752.

39. Chung, C.H.; Golub, E.E.; Forbes, E.; Tokuoka, T.; Shapiro, I.M. Mechanism of action of beta-glycerophosphate on bone cell mineralization. Calcif Tissue Int 1992, 51, 305–311, doi:10.1007/BF00334492.

40. Hadzir, S.N.; Ibrahim, S.N.; Abdul Wahab, R.M.; Zainol Abidin, I.Z.; Senafi, S.; Ariffin, Z.Z.; Abdul Razak, M.; Zainal Ariffin, S.H. Ascorbic acid induces osteoblast differentiation of human suspension mononuclear cells. Cytotherapy 2014, 16, 674–682, doi:10.1016/j.jcyt.2013.07.013.

41. Franceschi, R.T.; Iyer, B.S.; Cui, Y. Effects of ascorbic acid on collagen matrix formation and osteoblast differentiation in murine MC3T3-E1 cells. J Bone Miner Res 1994, 9, 843–854, doi:10.1002/jbmr.5650090610.

42. Carinci, F.; Pezzetti, F.; Spina, A.M.; Palmieri, A.; Laino, G.; De Rosa, A.; Farina, E.; Illiano, F.; Stabellini, G.; Perrotti, V., et al. Effect of Vitamin C on pre-osteoblast gene expression. Arch Oral Biol 2005, 50, 481–496, doi:10.1016/j.archoralbio.2004.11.006.

43. Xing, W.; Pourteymoor, S.; Mohan, S. Ascorbic acid regulates osterix expression in osteoblasts by activation of prolyl hydroxylase and ubiquitination-mediated proteosomal degradation pathway. Physiol Genomics 2011, 43, 749–757, doi:10.1152/physiolgenomics.00229.2010.

44. Suzuki, H.; Nezaki, Y.; Kuno, E.; Sugiyama, I.; Mizutani, A.; Tsukagoshi, N. Functional roles of the tissue inhibitor of metalloproteinase 3 (TIMP-3) during ascorbate-induced differentiation of osteoblastic MC3T3-E1 cells. Biosci Biotechnol Biochem 2003, 67, 1737–1743, doi:10.1271/bbb.67.1737.

45. Takamizawa, S.; Maehata, Y.; Imai, K.; Senoo, H.; Sato, S.; Hata, R. Effects of ascorbic acid and ascorbic acid 2-phosphate, a long-acting vitamin C derivative, on the proliferation and differentiation of human osteoblast-like cells. Cell Biol Int 2004, 28, 255–265, doi:10.1016/j.cellbi.2004.01.010.

46. Choi, J.Y.; Lee, B.H.; Song, K.B.; Park, R.W.; Kim, I.S.; Sohn, K.Y.; Jo, J.S.; Ryoo, H.M. Expression patterns of bone-related proteins during osteoblastic differentiation in MC3T3-E1 cells. J Cell Biochem 1996, 61, 609–618, doi:10.1002/(SICI)1097-4644(19960616)61:4%3C609::AID-JCB15%3E3.0.CO;2-A.

47. Zoch, M.L.; Clemens, T.L.; Riddle, R.C. New insights into the biology of osteocalcin. Bone 2016, 82, 42–49, doi:10.1016/j.bone.2015.05.046.

48. Golub, E.E.; Harrison, G.; Taylor, A.G.; Camper, S.; Shapiro, I.M. The role of alkaline phosphatase in cartilage mineralization. Bone Miner 1992, 17, 273–278, doi:10.1016/0169-6009(92)90750-8.

49. Ahmad Hairi, H.; Jayusman, P.A.; Shuid, A.N. Revisiting Resveratrol as an Osteoprotective Agent: Molecular Evidence from In Vivo and In Vitro Studies. Biomedicines 2023, 11, doi:10.3390/biomedicines11051453.

50. Iantomasi, T.; Romagnoli, C.; Palmini, G.; Donati, S.; Falsetti, I.; Miglietta, F.; Aurilia, C.; Marini, F.; Giusti, F.; Brandi, M.L. Oxidative Stress and Inflammation in Osteoporosis: Molecular Mechanisms Involved and the Relationship with microRNAs. Int J Mol Sci 2023, 24, doi:10.3390/ijms24043772.

51. Zhivodernikov, I.V.; Kirichenko, T.V.; Markina, Y.V.; Postnov, A.Y.; Markin, A.M. Molecular and Cellular Mechanisms of Osteoporosis. Int J Mol Sci 2023, 24, doi:10.3390/ijms242115772.

52. Rendina-Ruedy, E.; Rosen, C.J. Parathyroid hormone (PTH) regulation of metabolic homeostasis: An old dog teaches us new tricks. Mol Metab 2022, 60, 101480, doi:10.1016/j.molmet.2022.101480.

53. Yan, C.; Shi, Y.; Yuan, L.; Lv, D.; Sun, B.; Wang, J.; Liu, X.; An, F. Mitochondrial quality control and its role in osteoporosis. Front Endocrinol (Lausanne) 2023, 14, 1077058, doi:10.3389/fendo.2023.1077058.

54. Sabini, E.; Arboit, L.; Khan, M.P.; Lanzolla, G.; Schipani, E. Oxidative phosphorylation in bone cells. Bone Rep 2023, 18, 101688, doi:10.1016/j.bonr.2023.101688.

55. Yan, Q.; Jiang, J.; Xie, J.; Jiang, S.; Bai, Y. Inhibition of osteosarcoma cell proliferation in vitro and tumor growth in vivo in mice model by alstonine through AMPK-activation and PGC-1alpha/TFAM up-regulation. Acta Biochim Pol 2022, 69, 543–549, doi:10.18388/abp.2020_5769.

56. An, J.H.; Yang, J.Y.; Ahn, B.Y.; Cho, S.W.; Jung, J.Y.; Cho, H.Y.; Cho, Y.M.; Kim, S.W.; Park, K.S.; Kim, S.Y., et al. Enhanced mitochondrial biogenesis contributes to Wnt induced osteoblastic differentiation of C3H10T1/2 cells. Bone 2010, 47, 140–150, doi:10.1016/j.bone.2010.04.593.

57. Fan, Z.Q.; Bai, S.C.; Xu, Q.; Li, Z.J.; Cui, W.H.; Li, H.; Li, X.H.; Zhang, H.F. Oxidative Stress Induced Osteocyte Apoptosis in Steroid-Induced Femoral Head Necrosis. Orthop Surg 2021, 13, 2145–2152, doi:10.1111/os.13127.

58. Bai, S.C.; Xu, Q.; Li, H.; Qin, Y.F.; Song, L.C.; Wang, C.G.; Cui, W.H.; Zheng, Z.; Yan, D.W.; Li, Z.J., et al. NADPH Oxidase Isoforms Are Involved in Glucocorticoid-Induced Preosteoblast Apoptosis. Oxid Med Cell Longev 2019, 2019, 9192413, doi:10.1155/2019/9192413.

