## Supplementary material for "Defining the most potent osteoinductive culture conditions for MC3T3-E1 cells reveals no implication of oxidative stress or energy metabolism": Table S1

| **Gene** | **Primer sequences** |
| --- | --- |
| ***Alpl*** | Forward: 5’-CCTTGAAAAATGCCCTGAAA-3’  Reverse: 5’-CTTGGAGAGAGCCACAAAGG-3’ |
| ***Col1a1*** | Forward: 5’-TGGCCCCATTGGTAACGTTGGT-3’  Reverse: 5’-AGGACCTTGTTTGCCGGGTTCA-3’ |
| ***Runx2*** | Forward: 5’-TCTGCCGAGCTACGAAATGCCT-3’  Reverse: 5’-TGAAACTCTTGCCTCGTCCGCT-3’ |
| ***Bglap*** | Forward: 5’-CATGAGGACCCTCTCTCTGC-3’  Reverse: 5’-TGGACATGAAGGCTTTGTCA-3’ |
| ***Pgc1α*** | Forward: 5’-gacaggtgccttcagttcac-3’  Reverse: 5’-caaccagagcagcacactcta-3’ |
| ***Sod2*** | Forward: 5’-ttaacgcgcagatcatgca-3’  Reverse: 5’-ggtggcgttgagattgttca-3’ |
| ***Tfam*** | Forward: 5’-cataggcaccgtattgcgtg-3’  Reverse: 5’-tcggaatacagacaagactgataga-3’ |
| ***Actb*** | Forward: 5’-GCCCTGAGGCTCTTTTCCAG-3’  Reverse: 5’-TGCCACAGGATTCCATACCC-3’ |

**Table S1:** Sequences of the primers used for qPCR.
